# The Impact of Multivesicular Release on the Transmission of Sensory Information by Ribbon Synapses

**DOI:** 10.1101/2021.11.04.467256

**Authors:** Ben James, Pawel Piekarz, José Moya-Diaz, Leon Lagnado

## Abstract

The statistics of vesicle release determine how information is transferred at the synapse but the classical Poisson model of independent fusion does not hold at the first stages of vision and hearing. There, ribbon synapses encoding analogue signals can coordinate the release of two or more vesicles as a single event. The implications of such multivesicular release (MVR) for spike generation are not known. Here we investigate the hypothesis that MVR might provide advantages over Poisson synapses. We used leaky integrate-and-fire models incorporating the statistics of release measured experimentally from glutamatergic synapses of retinal bipolar cells and compared these with models assuming Poisson inputs constrained to operate at the same average rates. We find that MVR can increase the number of spikes generated per vesicle while reducing interspike intervals and latency to first spike. The combined effect was to increase the efficiency of information transfer (bits per spike) over a range of conditions mimicing retinal ganglion cells of different size. MVR was most advantageous in neurons with short time-constants and reliable synaptic inputs, when a lower degree of convergence was required to trigger spikes. In the special case of a single input driving a neuron with short time-constant, as occurs at the first synapse in hearing, MVR increased information transfer whenever spike generation required more than one vesicle. This study demonstrates how presynaptic integration of vesicles by MVR has the potential to compensate for less effective summation post-synaptically to increase the efficiency with which sensory information is transmitted.

## Introduction

The encoding of sensory information by spikes has been studied in detail (Gollisch and Meister, 2008; Kayser et al., 2009) but less is understood about the vesicle code that drives spiking. This is especially true in the early stages of vision and hearing where analogue voltage signals are transmitted through ribbon-type synapses (Lagnado and Schmitz, 2015; Neef et al., 2007). It has often been assumed that ribbon synapses recode graded signals by modulating the mean rate of a Poisson process in which vesicles fuse independently (Choi et al., 2005; Odermatt et al., 2012; Sterling and Laughlin, 2015) but an accumulation of evidence now indicates that the release of two or more vesicles can be tightly synchronized (Glowatzki and Fuchs, 2002; Hays et al., 2021; James et al., 2019; Singer et al., 2004). In retinal bipolar cells, for instance, the ratio of the variance to the mean of the exocytic response to a repeated stimulus is not one, as expected for a Poisson process, but about three (Moya-Díaz et al., 2022). Further, the fusion of multiple vesicles can be synchronized to within the resolution of electrophysiological recordings (<100 µs) by a process that has been termed *coordinated* multivesicular release to emphasize the lack of independence (Singer et al., 2004). At central synapses the release of two or more vesicles within milliseconds is also termed multivesicular release (MVR) (Auger et al., 1998; Christie and Jahr, 2006; Holler et al., 2021; Huang et al., 2010) but it may be that in these cases it simply reflects a large increase in release probability.

Whatever the mechanisms underlying MVR, the closer together in time vesicles are released at a single synaptic connection the more effectively they will summate on a postsynaptic dendrite to impact the spiking activity of the target neuron (Li and Ascoli, 2008). This simple idea has profound implications for understanding how information is transmitted in the brain because in principle it might allow for a sensory stimulus to be represented not just by the frequency of synaptic events (a temporal code) but also by their *amplitude* (Holler et al., 2021; James et al., 2019; Lisman et al., 2007). Direct evidence for amplitude coding at a synapse has been provided by improvements in the optical imaging of glutamate in the retina, which provide the resolution to count vesicles as they are released from individual active zones. This approach reveals that bipolar cells transmit visual information both in the rate of vesicle release and in the number of vesicles released within each event, with larger and rarer events carrying the most information per vesicle (James et al., 2019).

We now need to understand how coordinated MVR contributes to the conversion of an analogue sensory signal into spikes. An experimental investigation of this question is difficult because it would require spikes to be recorded post-synaptically while also characterizing the influence of individual active zones monitored with a resolution of single vesicles. We have therefore started with a modelling approach in which a train of synaptic events is simulated based on experimentally measured statistics of vesicle release and used as the input to a Leaky Integrate-and-Fire (LIF) model of the post-synaptic neuron (Abbott, 1999; Burkitt, 2006a). We gather these statistics from the ribbon-type synapse of retinal bipolar cells across which visual information is transmitted to post-synaptic ganglion cells and then focus on the special case of transmission of auditory information from the ribbon synapses of hair cells. A comparison with simulations in which synapses release vesicles independently demonstrates that MVR can increase the efficiency with vesicles are used to transmit sensory information over a range of conditions mimicking retinal ganglion cells of different sizes. In the special case of auditory hair cells driving afferent fibers through a single synapse, MVR increased the efficiency of information transfer whenever spike generation required depolarization greater than that caused by a single vesicle.

## Methods

### Modelling Overview

Modelling the effects of MVR on spike generation involved the following basic steps:

#### 1. Statistics of vesicle release in retinal bipolar cells responding to visual stimuli

Using iGluSnFR expressed in retinal bipolar cells of larval zebrafish, we measured the amplitude of synaptic events and their timing relative to a sinusoidal full-field stimulus of varying contrast (Fig. 1A). At each contrast we calculated i) the average rate of vesicle release over a 30 s period *R* (Fig. 1B; n = 60 synapses); ii) the probability of events composed of different numbers of quanta, *PQe* (Fig. 1C; n = 55 synapses), and iii) the “temporal jitter” of events of each amplitude, quantified as the standard deviation of event times relative to any particular phase of the sinusoidal stimulus (*TJ(Q)*, Fig. 1D-F; n = 60 synapses). Each statistic was calculated from at least 20 zebrafish.

**Figure 1.**
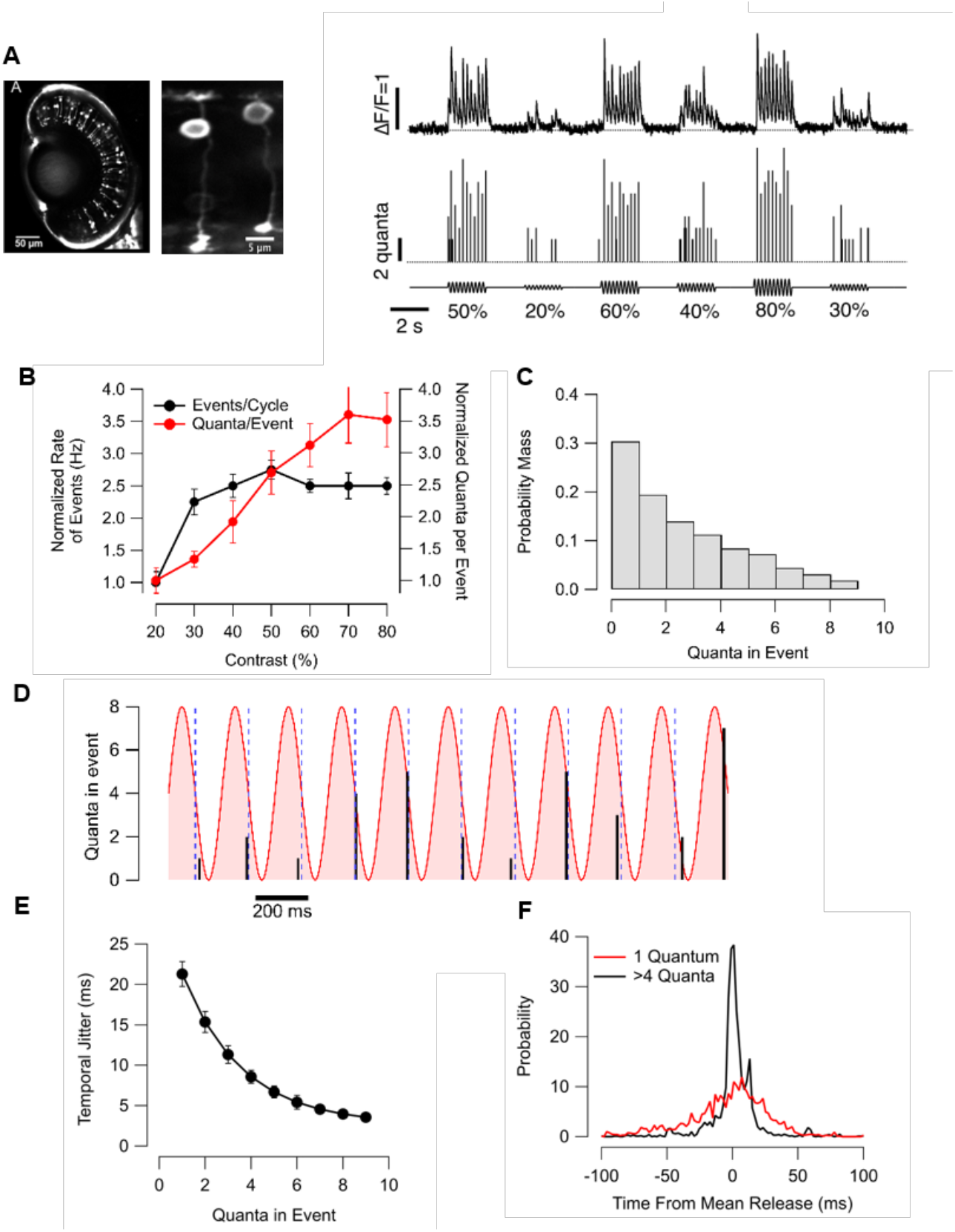
Multivesicular Release in the Retina of Zebrafish. **A**. Left: Multiphoton image of BCs in the retina of a larval zebrafish. They are visualized by the iGluSnFR reporter in the surface membrane. Middle: Two BCs at higher magnification. The mosaic expression of the reporter allows for distinguishing individual BCs and distinguishing glutamate release at different synapses. Right Top: Recorded fluorescence time series from an individual active zone. Bottom: Estimated quantal content of the time series above. Note the heterogeneity in the response amplitude, corresponding to variable quanta released per event. **B**. Normalized change in the rate of events (black) and average event amplitude in quanta (red) (n = 60 synapses). **C**. Distribution of event amplitudes during a 30 s application of a 5 Hz stimulus at 80% contrast (n = 55 synapses), truncated to only include events containing fewer than 11 quanta. **D**. A quantal time series (black lines) in response to a 5 Hz, 80% contrast full-field stimulus (red), with blue dashed lines present only for ease of visibility. Note that events composed of four or more vesicles occur in a temporally small window during each cycle, while lower quantal events are more temporally dispersed. **E**. The temporal jitter as a function of quanta in event for a 100% contrast stimulus (n = 60 synapses). **F**. Mean-shifted distribution of events containing 5 or more quanta (black) and of events containing one quantum (red) in response to a 30 s, 5 Hz, 100% contrast sinusoidal stimuli. Data from E.

#### 2. Simulation of synapse output

Based on step 1, two types of simulated synaptic output were constructed: one in which stimulus contrast was represented as a “pure” rate-code in which all vesicles were released independently and a second “hybrid code” in which amplitude also varied (Fig. 2A). In both cases, the overall average rate of vesicle release *R* was fixed to values measured experimentally at a given contrast. The number of vesicles in an event (for the MVR case) or the number of released uniquantal events (for the rate case) was sampled from the distribution PQ. For the MVR case, each event was then given a time by sampling from the Gaussian distribution of event times with mean and variance measured experimentally i.e by TJ(q) shown in Fig. 1F. For the rate case, a corresponding number of event times were sampled from the distribution of uniquantal event times.

**Figure 2.**
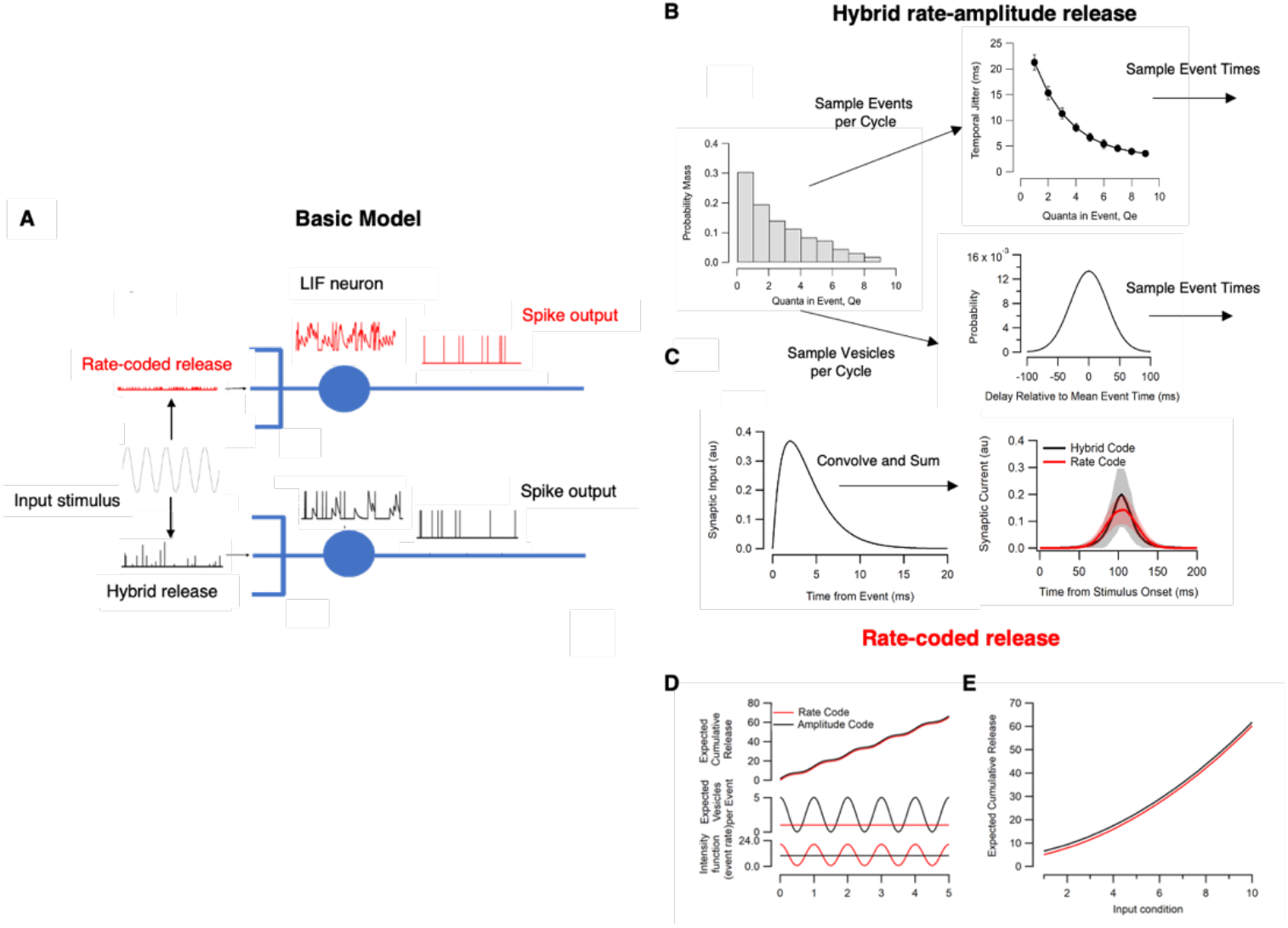
Overview of modelling. **A**. A leaky-integrate-and-fire neuron received either rate (top) or hybrid (bottom) inputs. **B**. Illustration of release simulation. The number of vesicles in an event (for the MVR case) or the number of released uniquantal events (for the rate case) is sampled from the distribution PQ. For the MVR case, each event is then given a time by sampling from a Gaussian Distribution with variances set by the mean temporal precision of events, given by TJ(q). For the rate case, a corresponding number of event times are sampled from the distribution of uniquantal event times (bottom middle). **C**. The sampled event time and amplitude tuples are then convolved with a synaptic current filter. The resulting current is summed over the number of inputs to the cell. Plotted on the right are the average synaptic current input to a cell utilizing a rate code (red) and hybrid code (black). Note the increased temporal variability in the rate code (SD = 360 ms for the hybrid case vs. 685 ms for the rate case). **D**. Demonstration of the Poisson release model. Bottom: intensity function driving the rate of vesicular events. Middle: expected number of vesicles released in an event as a function of time. Top: Expected cumulative vesicles released as a function of time for rate and amplitude codes, offset for visualization. These functions have been equalized by design. **E**. Expected total vesicles released over a one second window as a function of stimulus ‘contrast’.

The output of a synapse employing coordinated MVR was generated by randomly selecting a total of N events with amplitudes chosen from the distribution of quanta in an event P(Qe) (Fig. 2B). Each event was then given a time by sampling from the Gaussian distribution of event times with mean and variance measured experimentally i.e by TJ(Qe) shown in Fig. 1F. Each synapse was constrained to generate only one or zero events per cycle of the stimulus, with the probability of no event given by *p*_*Q*_(*f, c*). The output of each synapse was therefore described as a vector E of event times and a vector Qe of event amplitudes, both of length N.

The rate-code comparison was constructed by replacing each event consisting of more than one quantum with an equivalent number of uni-quantal events, each event time now sampled from the distribution of times of uniquantal events (Fig. 1F). The vectors describing a pure rate synapse were therefore of variable lengths corresponding to the total number of vesicles sampled within the 30 s sample period. The resulting vectors Qe and E of event quanta and event times were then passed into a conductance-based model LIF model (Fig. 1) and the spike output analysed.

#### 3. Simulation of synapse output using marked Poisson Processes

For purely rate-coded synapses, we used a Nonhomogeneous Poisson Process (NHPP) as the driving force for vesicle release, in which the mean rate λ(t) varied sinusoidally in time. By definition, simple Poisson Processes describe a situation in which all events occur independently and with exponentially distributed interevent times and cannot, therefore, be used to model MVR when two or more vesicles are released within a single synaptic event. To construct an amplitude-modulated Poisson Process we turned to a modification known as Poisson Splitting or Location-dependent thinning (Resnick, 1992), where events can be assigned a type based upon their location in time. In this case events occur with a constant rate but the number of vesicles in each event is chosen based upon when the event occurred. We could therefore drive the LIF model neuron in two distinct regimes: a pure rate case, where all information is encoded by modulating the mean rate of release of individual vesicles and a hybrid case, where information is encoded by modulating both the rate and quantal content of synaptic events.

A key aspect of our investigation was to construct the rate and hybrid models in such a way that each released the same average number of vesicles, with differences found only in the variabilities. In other words, vesicles were released individually in the “rate code”, but in packets of variable size in the “hybrid code”. To simulate pure rate-coded synaptic inputs case, we used a sinusoidal intensity function. The response of each synapse to varying contrasts was simulated by increasing the amplitude of the sinusoid between a total release of between 5 vesicles s^-1^ at input condition 1 (which we term 10% contrast below) and 65 vesicles s^-1^ at input condition 10 (100% contrast; Fig. 2D and E). for the rate case. Responses from multiple cells were assumed to come from the same distribution, and as such we simulated the responses of multiple BCs onto a single RGC by summing the response to all the individual synaptic inputs. To simulate hybrid coding the probability of release was also modulated as a sinusoid with an amplitude generating an average rate of vesicle corresponding to that measured experimentally at a given stimulus contrast (Fig. 1A).

#### 4. A conductance-based Leaky Integrate-and-Fire (LIF) model of the post-synaptic neuron

The vectors describing amplitudes and times of synaptic events were convolved with a synaptic filter to estimate the time-course of the synaptic current Sin (Fig. 2C) which was then passed through a Leaky Integrate-and-Fire model to calculate the change in voltage and spike output. LIF models provide a computationally compact method for exploring basic physiological parameters affecting spiking, including the number of synaptic inputs, cell time-constant, threshold relative to resting potential and ion channel conductances (Abbott, 1999; Burkitt, 2006a, b; Gerstner et al., 1997). Voltage is governed by the differential equation:

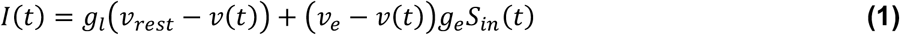

Where *g*_*l*_ is leak conductance, *g*_*e*_ is excitatory conductance, *v*_*e*_ is the excitatory reversal potential, and *v*_r*e*st_ is the resting membrane potential. The synaptic input, *S*_*in*_, is defined by the convolution of an alpha current impulse function with the input s(t),

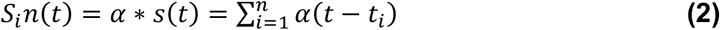

Where s(t) is the sequence of Dirac delta function describing the timing of each event in a realization of the Process, and

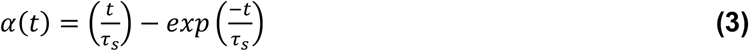

describes the time course of synaptic current. Here, we used a value of τ_s_ = 2 ms (Destexhe and Rudolph, 2004) which yields a total conductance of 2 nS per quanta. Once the cell hits a threshold θ, it fires a spike and the voltage is immediately reset to the reset potential v_*reset*_, and remains at that value in a refractory period for τ_r*e*f_ = 5 ms. There is no explicit membrane time-constant (τ_m_ = RC) in a conductance model but the time course of the leak varies as the reciprocal of the leak conductance 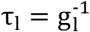.Due to its similarity with the current-based model, we will describe the model in terms of τ_*l*._ The ratio between the membrane capacitance and the leak channel conductance is the membrane time-constant 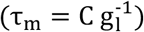.

To compare how the MVR might shape the spike output of a postsynaptic cell across a range of physiological values, we systematically varied the parameters shown in Table

1. Note that here, rather than representing the set of the excitatory conductances, g_*e*_, we computed the approximate leak conductance required for the cell to spike in response to k vesicles in a perfect integrator – with corresponding *g*_*e*_ values ranging in the 1-100 nS range. In the retina, bipolar cells transmit to RGCs of widely varying size and electrical properties. In mice, for instance, input resistances of RGCs can vary from dozens of MΩ to a GΩ, with values of the membrane time-constant ranging from less than 5 ms to over 50 ms (O’Brien et al., 2002). At another extreme, primary auditory afferents receive inputs from single active zones delivering the output from hair cells and have input resistances of GΩ leading to the suggestion that a single glutamatergic vesicle is sufficient to trigger a spike (Fuchs and Glowatzki, 2015; Glowatzki and Fuchs, 2002). Thus, the number of vesicles sufficient to generate a spike in a cell postsynaptic to a ribbon synapse (k) varies from one to hundreds.

#### 5. Analyzing the spike output of the model neuron

We concentrated on three aspects of the spiking response of the model neuron: the spike count generated by each cycle of the 5 Hz stimulus, the temporal accuracy of spikes and the average rate of information transfer about the modulation in stimulus strength (i.e contrast).

Temporal accuracy was quantified as “temporal jitter” (TJ, in ms), which was in turn calculated from the Vector Strength (VS), which takes values between zero and one, with zero being complete independence from the stimulus and one being perfect phase-locking (Baden et al., 2011; Goldberg and Brown, 1969). VS was computed for each quantal event type in response to a 30 s stimulus of varying contrast delivered at 5 Hz, with the equation:

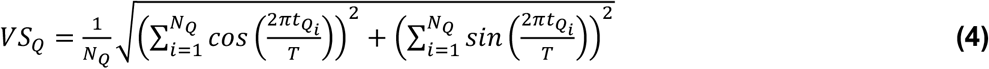

where 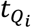 is the phase of the i^th^ event of quantal content Q, and T is the stimulus period. VS can then be easily converted to Temporal Jitter (TJ) by:

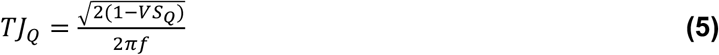

We calculated the mutual information (Shannon, 1948) between the stimulus S and either the spike times or the total number of spikes occurring over a period of 200 ms (N, corresponding to one cycle of the 5 Hz stimulus, 2 cycles of the 10 Hz stimulus or 3 cycles of the 15 Hz stimulus). Mutual information for spike counts was computed empirically from the joint and marginal distributions *p*(*s, n*), *p*(*n*), and *p*(*s*) as:

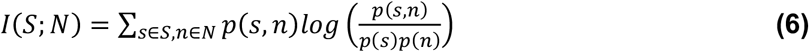

using the direct approach (Victor, 2006). Each stimulus *s* is has one of three contrasts and one of three frequencies, leading to a total of 9 different stimuli and a maximum of ≈ 3.17 bits of information. The total mutual information was calculated by discretising the response in a 200 ms time window into 5 ms bins, yielding a total of 40 response bins and 2^40^ possible responses (the vast majority of which are never observed). To reduce bias in the estimates, each simulation was run a total of 1000 times, with typical simulations generating under 80 unique responses. Keeping the ratio of the number of samples to the number of distinct responses large ensures any bias is small (Panzeri et al., 2007). The spike count information was computed similarly, using the distribution of spike counts **N**. Finally, the spike-time information was computed as the difference between the total information and the spike count information, *I*(*S; R*)_*time*_ = *I*(*S; R*)_*Total*_ − *I*(*S; R*)_*Count*_

#### The signal-to-noise ratio associated with a change in the rate of a Poisson process

Imagine the mean rate of a Poisson process, R, changes by a factor a. The signal, S, generated by comparing two observation times Dt will be the change in the mean number of events counted in each period (aR Dt - R Dt), and the variance in that signal will be the sum of the number of events counted in each period (aR Dt + R Dt). Defining the SNR in the same way as the discriminability (d’) used in signal detection theory, we have

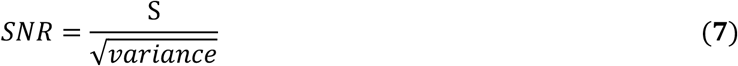

yields

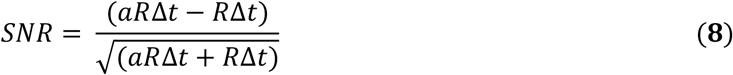

which can be rearranged to obtain the time period Dt required to obtain a given SNR, as

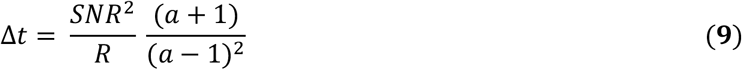

#### Multiphoton imaging of glutamate release

Methods for *in vivo* imaging of glutamate release in the retina of zebrafish were as described previously (James et al., 2019). Transgenic larval zebrafish expressing the iGluSnFR reporter under the *ribeye a* promoter were imaged at 7-9 days post-fertilization after embedding in a 3% low melting-point agarose in E2 on a glass cover-slip. To prevent eye movements, the fish was injected with 1 nL of alpha-bungarotoxin behind the eye. Imaging was carried out using a multiphoton microscope (Scientifica) equipped with a mode-locked titanium–sapphire laser (Chameleon, Coherent) tuned to 915 nm and an Olympus XLUMPlanFI ×20 water immersion objective (NA of 0.95). Fluorescence emission was also collected using a sub-stage oil condenser (Olympus; NA of 1.4), which was essential to achieve a SNR adequate for reliable detection of single vesicles. A total of 42 zebrafish were used in this study and all measuremnts were made in the afternoon.

Light emission was filtered through GFP filters (HQe525/50, Chroma Technology) before detection using GaAsP photomultipliers (H7422P-40, Hamamatsu). Photocurrents were passed to a current-to-voltage converter, after which signals from the objective and condenser were summed and digitized through ScanImage (v.3.6; Vidrio Technologies). For this work, all time-series were acquired as 128 pixel line scans imaged at 1 kHz and analysed in custom software written within IgorPro (Wavemetrics). Stimuli were generated by an amber LED (peak wavelength of 590 nm) filtered through 590/10 bandpass filter (Thorlabs) delivered through a light-guide placed near the fish’s eye, with the mean stimulus intensity of ∼320 nW mm^−2^. Detailed evidence that this approach can detect the release of single vesicles from individual active zones has been presented previously (James et al., 2019).

## Results

### Synaptic coding of a sensory stimulus: gathering release statistics for an empirically-driven model

Our aim was to investigate how the statistical properties of vesicle release affected the synaptic transfer of sensory information, comparing synapses employing coordinated MVR with the conventional view of synapses operating as simple Poisson machines. The starting point was to gather experimental measurements of release statistics from retinal bipolar cells expressing the glutamate reporter iGluSnFR, and examples of such records are shown in Fig. 1A. Both the rate and amplitude of glutamate transients varied widely during sinusoidal modulations in light intensity (James et al., 2019).

We measured how changes in contrast altered three aspects of the vesicle code highlighted in Fig 1: event rate R(Q), distribution of event amplitudes P(Qe), and variability of event timing TJ(Q). R saturated at lower contrasts than Qe, such that contrasts greater than ∼50% were often encoded entirely by the number of vesicles released within an event (Fig. 1B and C). A potential role of MVR in determining the temporal accuracy of spiking in the post-synaptic neuron is immediately suggested by the observation that multiquantal events were more tightly phase-locked to the stimulus than vesicles released individually (Fig. 1D-F). Uniquantal events had a temporal jitter of 21.2 ± 1.5 ms, while events composed of 9 quanta had a temporal jitter of just 2.7 ± 0.6 ms, which is very similar to the precision of spikes observed in RGCs of the salamander retina responding to a high-contrast stimulus (Berry et al., 1997a; Uzzell and Chichilnisky, 2004). The relation between TJ and Qe is shown in Fig. 1E and a comparison of the distribution of times for uniquantal and multiquantal events is shown in Fig. 1F.

Based on these measurements we constructed two types of synaptic input to the model neuron: purely rate-code inputs in which all vesicles were released independently (red in Fig. 2A) and hybrid inputs in which both rate and amplitude of events varied (black; see Methods). To compare how these two coding strategies performed with equivalent resources the average rates of vesicle release were fixed for both types of input at the values measured experimentally at a given contrast and frequency (Fig. 1B). Measuring the output of individual synapses to stimuli of different contrasts then allowed us to make measures of the mutual information between the stimulus set and spikes generated in model neurons.

### Synaptic coding strategy alters interspike intervals

Examples of simulations comparing the effects of rate-coded and hybrid-coded synaptic inputs are shown in Fig. 3 for two model neurons chosen to highlight the effect of cell size. Bipolar cells transmit the visual signal to RGCs that vary in the dimensions of their dendritic trees (determining the size of receptive fields) and membrane time-constants (determining temporal integration of synaptic inputs). In the fovea of primates, for instance, midget ganglion cells have small receptive fields and are electrically compact, receiving 25-50 synaptic inputs from between one and three bipolar cells (Calkins et al., 1994; Kolb and Marshak, 2003). But in the periphery of the retina, RGCs have larger dendritic fields and receive inputs from hundreds of bipolar cells, with up to thousands of synapses (Jacoby et al., 2000). We therefore begin by comparing spike outputs in a “small” synthetic RGC (τ_*l*_ = 50 ms with 50 synaptic inputs) and a “big” RGC (τ_*l*_ = 5 ms with 500 inputs) to stimuli that are spatially uniform, acting on all excitatory inputs equally. These two example neurons have different contrast-response relation (Fig. 3A), qualitatively similar to variations observed physiologically.

**Figure 3.**
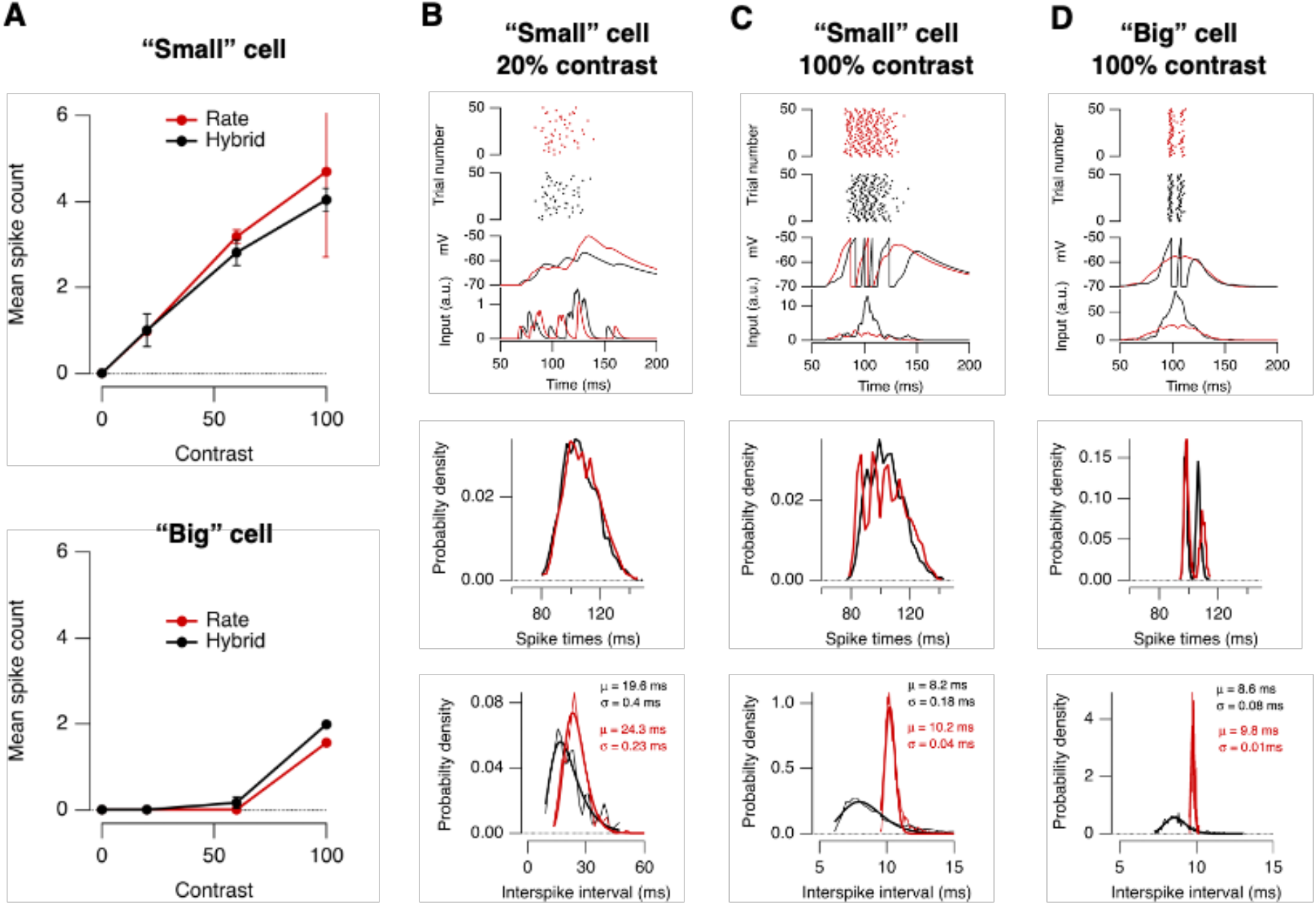
Comparison of vesicle codes on spike generation in a “small” cell and a “large” cell. **A**. Mean spike count as a function of contrast for a “small” cell (top) and “big” cell (bottom). The “small” cell was simulated with: inputs = 50, k = 10, τ_*l*_ = 50 ms, *v*_*e*_ = 0, *θ* = −50 mV, vr_*e*st_ = −70 mV. The “big” cell was simulated with: inputs = 500, k = 100, τ_*l*_ = 5 ms, *v*_*e*_ = 0, *θ* = −50 mV, _r*e*st_ = −70 mV. Stimuus at 5 Hz. Red shows responses to synaptic inputs employing a pure rate code and black to inputs employing a hybrid rate-amplitude code. **B**. A small cell responding to 20% contrast. Top panel: raster plots of spike times (top), example voltage traces (middle), and example synaptic input (bottom), for hybrid and rate codes. One cycle of the sinusoidal stimulus is shown. Middle panel: distribution of spike times. Bottom panel: distribution of interspike intervals (ISI, thin trace) could be described as a lognormal function (bold trace) with values of µ and s shown (equation 10). **C**. Same as B, but for 100% contrast. Note the lower mean but broader distribution of ISI’s in the cell receiving hybrid inputs. **D**. Same as C, but for the big cell. Note that despite the much narrower distributions of spike times (middle) the ISI’s were more broadly distributed in the cell receiving hybrid inputs (bottom).

Using a stimulus of low contrast (20%), the big cell barely responded but the small cell generated an average of one spike per cycle of the 5 Hz stimulus (Fig. 3A). The distributions of spike times were not significantly affected by the coding strategy of the synaptic inputs (Fig. 3B, middle) but when two or more spikes were generated within a cycle the distribution of interspike intervals (ISI) was shifted to significantly lower values when input was provided by hybrid synapses employing MVR (Fig. 3B, bottom; p < 0.02; Kolmogorov–Smirnov test). This distribution, Y(ISI), could be described as a lognormal function of the form

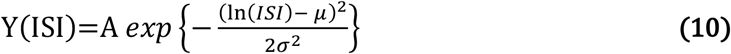

where µ and σ are the mean and standard deviation of a normal distribution in ln(ISI). A lognormal distribution of ISI’s is commonly found in RGCs (Levine, 1991) and was a consistent feature of the spike output across a range of conditions (Fig. 3B-D). In the small cell responding to 20% contrast, the ISI was reduced from 24.3 ± 0.2 ms with rate-coded inputs to 19.6 ± 0.4 ms with hybrid inputs. Stimulating the small cell with 100% contrast increased the average spike count by a factor of more than 4 (Fig. 3A) but the same qualitative difference was observed (Fig. 3C, bottom): the mean ISI was significantly smaller with hybrid inputs (8.2 ± 0.18 ms) compared to rate-coded inputs (10.2 ± 0.04 ms; p < 0.0001, KS test). The big cell with ten times as many inputs and one-tenth the time-constant generated about half as many spikes at 100% contrast, but again the mean ISI was shorter with hybrid inputs (Fig.3D, bottom; p < 10^−20^; KS test).

The precise timing of spikes in an RGC can code information about the stimulus (Ishii and Hosoya, 2020; Meister and Berry, 1999) that neurons post-synaptic to RGCs can decode (Kaplan et al., 1987; Usrey and Reid, 1999). In mammals, for instance, recordings from RGCs and their target neurons in the lateral geniculate nucleus (LGN) show that the efficiency with which an RGC spike drives an LGN spike depends on the ISI: ISIs shorter than about 10 ms have the greatest efficiency but this decreases progressively with ISI up to ∼30 ms (Usrey et al., 1998). Such ISI-based filtering of retinal spikes can shape receptive fields and may serve as mechanism by which the information carried in ISI’s is decoded (Ishii and Hosoya, 2020; Rathbun et al., 2010). The simulations in Fig. 3 therefore show that a hybrid vesicle code driving excitation of RGCs has the potential to increase the visual information transmitted to post-synaptic targets by shortening spike intervals.

### Coordinated MVR increases spike count over a range of conditions

The spike response of RGCs will depend not only on the numbers of synaptic inputs (N) that converge on the dendritic tree but also on the strength of these inputs. In our modelling framework synaptic strength is directly related to k, the minimum number of vesicles that would need to be summed instantaneously to depolarise the neuron to threshold. Although we systematically varied both k and N (Table 1) we found the ratio k/N to be a useful summary metric that reduced dimensionality of the data while providing insight into the effects of both synapse strength and convergence. For instance, depolarization of the target neuron to threshold might occur by integrating signals from just a few synapses with a high probability of release (i.e reliable synapses, high k/N) or more synapses with lower average response (i.e synapses of lower reliability with low k/N). Medium-sized RGCs in guinea-pigs receive about 1000-2000 inputs, and a brief burst of excitation comprising 3-65 vesicles triggers a burst of 1-6 spikes (Freed, 2005). This example therefore provides an upper limit of k ∼10 while k/N is ∼0.01. In contrast, a midget RGC might receive just 10 inputs. If the same number of vesicles are required to trigger a spike, k/N = 1, and even if just one vesicle is sufficient, k/N =0.1.

**Table 1:** Simulation Parameters. Parameters used for simulations, with ranges based upon experimentally recorded values (Destexhe and Rudolph, 2004; O’Brien et al., 2002; Pillow et al., 2005)

| Parameter (symbol) | Simulated Value/s |
| --- | --- |
| Vesicles Required to Spike, $k$ | 10, 25, 50, 100 vesicles |
| Leak Constant, $\tau_l$ | 1, 5, 10, 20, 50 ms |
| Resting Potential, $v_{\text{rest}}$ | -70, -65, -50, -55 mV |
| Excitatory Reversal Potential, $v_e$ | 0 mV |
| Spiking Threshold, $\theta$ | -50, -45, -40, -35 mV |
| Number of Inputs, $N$ | 50, 100, 250, 500 |
| Refractory period, $\tau_{\text{ref}}$ | 5 ms |
| Synaptic Time Constant, $\tau_s$ | 2 ms |
| Stimulus Frequency, $f$ | 5, 10, 15 Hz |
| Stimulus Contrast, $c$ | 20, 60, 100% |

Examples of the interaction between k/N and the time-constant of the target neuron are shown in Fig. 4. Increasing τ_3_ always increased the spike count over a single stimulus cycle, and this is shown in Fig. 4A (top) for an example neuron with N = 500 inputs requiring the summation of K = 100 vesicles to trigger a spike (k/N = 0.2). The absolute difference in spike count between hybrid inputs and rate-coded inputs are shown below, demonstrating that for this value of k/N, the hybrid code produces more spikes than the rate code when τ_3_ is less than ∼15 ms. The interaction between τ_3_ and k/N over a range of values is shown for hybrid coded inputs in Fig. 4B and for rate-coded inputs in Fig. 4C. A clear pattern emerges: for either coding strategy, the lower the average number of vesicles released per synapse to trigger a spike the larger the total spike count for a cell of given time-constant. But the size of these changes were not the same and the difference between the counts generated by hybrid- and rate-coded inputs is shown in Fig. 4D, colour-coded such that increases are shown as depth of green and decreases as depth of red. For a given τ_3_, rate-coded inputs generate more spikes at lower k/N and hybrid inputs were more efficient at higher k/N. In other words, the rate code generated more spikes per vesicle than the hybrid code only when synapses were relatively unreliable. As the average number of vesicles released per synapse increased, a regime was entered in which the hybrid code generated spikes with greater efficiency, after which the coding strategy did not affect the spike count. The basic insight provided by these results is that where spike generation required more reliable synapses, it often became more effective if summation began *presynaptically* by the process of coordinated MVR.

**Figure 4.**
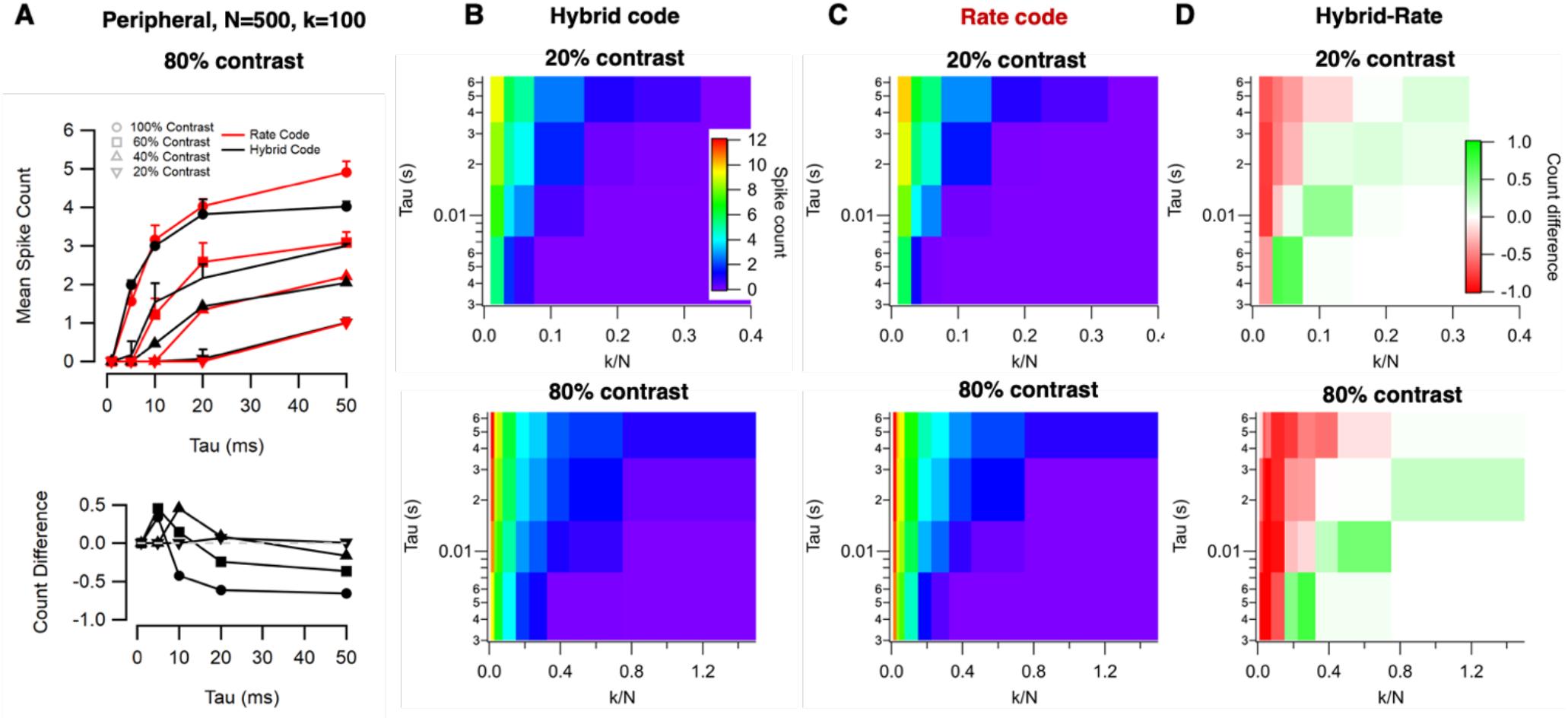
Hybrid coding increased spike count over a range of physiologically-relevant conditions. **A**. Top: Mean spike count as a function of leak time-constant (τ_l_) for a cell receiving N=500 inputs and requiring k=100 vesicles to spike in response to a 5 Hz stimulus of varying contrast (other parameters as in the “large peripheral” cell in Fig. 3). This count is over a 200 ms time window so can be multiplied by 5 to obtain spike rates. Bottom: Spike Count Difference (Hybrid – Rate) from the data in top panel. For this ratio of k:N, the hybrid code produces more spikes than the rate code when τ_l_ < 15 ms. **B**. Top: heat map showing the spike count generated by hybrid inputs for cells of varying k/N and τ_l_ for a stimulus of 20% contrast. Counts were averaged over values of k and N shown in Table 1. Bottom: as top but for 80% contrast. **C**. As B but for rate-coded inputs. Note again that the average rate of vesicle release was the same as in B for a given contrast. **D**. The difference between the counts generated by hybrid- and rate-coded inputs in B and C. Note that, for a given τ_l_, rate-coded inputs generate more spikes at lower k/N and hybrid inputs were more efficient at higher k/N.

How does this comparison of the hybrid and rate codes relate to the physiology of the retina? Compared to other visual circuits, such as those in the cortex, the retinal output is strikingly reproducible over repeated trials of a stimulus and contrasts of 20-40% can trigger spikes in RGCs of a variety of species with almost perfect reliability (i.e close to 100% of trials; (Berry et al., 1997a; Gollisch and Meister, 2008)). Unsurprisingly, the reliablity of individual bipolar cell synapses was lower. At 20% contrast a 5 Hz stimulus triggered a response in 31 ± 3% of stimulus cycles (N = 48 synapses), rising to 64 ± 4% of cycles at 60% contrast (N = 56 synapses). Experimental measurements of the average number of vesicles released per stimulus at these contrasts (where N is a single synapse) are shown in Fig. 1A-B: a stimulus of 20% contrast released an average of 2.32 ± 0.09 (N = 48 synapses) vesicles per synapse, while 60% contrast released 2.84 ± 0.14. Comparing these experimental measurements with the results of modelling indicates that synapses of bipolar cells are operating over a range of reliabilities where coordinated MVR provides an advantage over a pure rate-code by increasing the number of spikes generated per released vesicle.

### Coordinated MVR reduces spike latency over a range of conditions

Visual information is contained not only in the number of spikes generated by a stimulus but also in their timing (Butts et al., 2011; Gollisch and Meister, 2010), which can be precise to within a few milliseconds in RGCs (Berry et al., 1997a; Uzzell and Chichilnisky, 2004), the LGN (Reinagel and Reid, 2002) and visual cortex (Buracas et al., 1998). In RGCs, the temporal precision of spiking allows the latency to the first spike to transmit more information about a stimulus than the total number of spikes, at least when a stimulus drives episodic responses (Gollisch and Meister, 2008). To assess the impact of MVR on the temporal precision of spike generation we therefore analyzed how τ_3_ and k/N affecting first spike latencies (Fig. 5).

**Figure 5.**
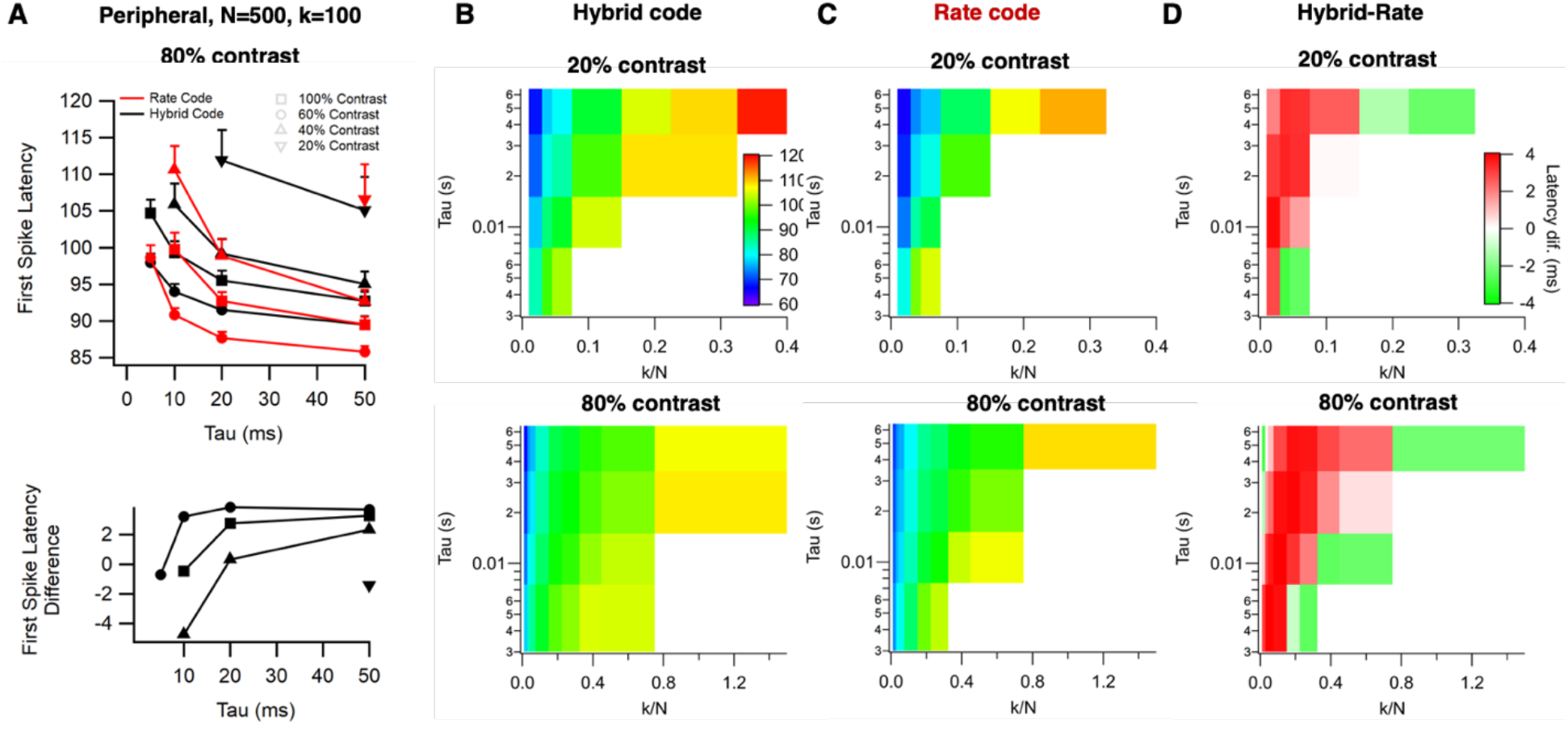
Hybrid coding reduced spike latency over a range of physiologically-relevant conditions. A. Top: Mean first spike latency as a function of leak time-constant (τ_l_) for a cell receiving N=500 inputs and requiring k=100 vesicles to spike in response to a 5 Hz stimulus of varying contrast (other parameters as in the “large peripheral” cell in Fig. 3). Latencies were measured relative to an arbitrarily chosen phase of the sinusoidal stimulus so it is the change in these values that is most informative. Bottom: Latency Difference (Hybrid – Rate) from the data in top panel. For this ratio of k:N, the hybrid code produced at greater latency when τ_l_ > 15 ms. B. Top: heat map showing the average first spike latency generated by hybrid inputs for cells of varying k/N and τ_l_ for a stimulus of 20% contrast. Latencies were averaged over values of k and N shown in Table 1. Bottom: as top but for 80% contrast. C. As B but for rate-coded inputs. D. The difference in first spike latency generated by hybrid- and rate-coded inputs in B and C. For a given τ_l_, rate-coded inputs generated faster responses at lower k/N while hybrid inputs generated faster responses at higher k/N (i.e when spikes are generated by activation of a larger proportion of inputs). Note the different ranges for k/N in upper and lower panels.

First spike latency depended on both the membrane time constant τ_3_ and the reliability of the synaptic inputs k/N (Fig. 5A-D). All but the very fastest time constants produced shorter latency spikes utilizing a rate code compared to a hybrid code, with delays being reduced by up to 4.5 ms. Cells requiring fewer vesicles to spike benefit from the wider temporal window of vesicle release in univesicular compared to multivesicular events (as seen in Fig. 1F). However, the more reliable the inputs the weaker this effect (Fig. 5D). At an intermediate range of k/N hybrid inputs generated spikes with shorter latencies and then at higher k/N there was no dependence on coding strategy. The general picture is that the rate code generates spikes with shorter latency only in larger neurons in which spiking is driven by larger numbers of less reliable synaptic inputs - conditions more likely to occur in brain regions such as the cortex than the retina or other circuits with more reliable synapses.

### Coordinated MVR increases information transmission across a range of conditions

Having established that MVR impacts both spike count and timing we asked how these effects interact to alter the transmission of information (Borst and Theunissen, 1999). The mutual information was calculated between a set of nine stimuli (contrasts of 20%, 60% and 100% at frequencies of 5 Hz, 10 Hz and 15 Hz) and each of three measures of the response in the model neuron: spike count, spike time and the total information within the spike sequence. A neuron that encodes a total of log2(9) = 3.17 bits of information over a 200 ms time window is discriminating perfectly between the nine stimuli.

The heat plots in Fig. 6 demonstrate that spike count carried more of the total information than spike time for both the synaptic coding strategies. But the relative advantages of MVR and rate codes again depended on the time-constant of the target neuron and the reliability of the inputs driving spikes. In a cell of given time-constant, an increase in the reliability of inputs caused the hybrid code to become more efficient at generating spikes (Fig. 4) with shorter latency (Fig. 5) resulting in more information transmission (Fig. 6). In the small cell, for instance, the spike count over a 200 ms period carried 0.07 bits more information when generated by hybrid inputs, while spike time carried 0.08 bits more, and the total information difference was ∼0.16 bits (equivalent to an information rate of 0.82 bits s^-1^). The largest difference in information rate in favour of the hybrid code was 3.8 bits s^-1^ or 24% of the maximum possible information rate, occurring when the post-synaptic cell required 25 vesicles to spike, had 250 inputs, and a 1 ms time constant.

**Figure 6.**
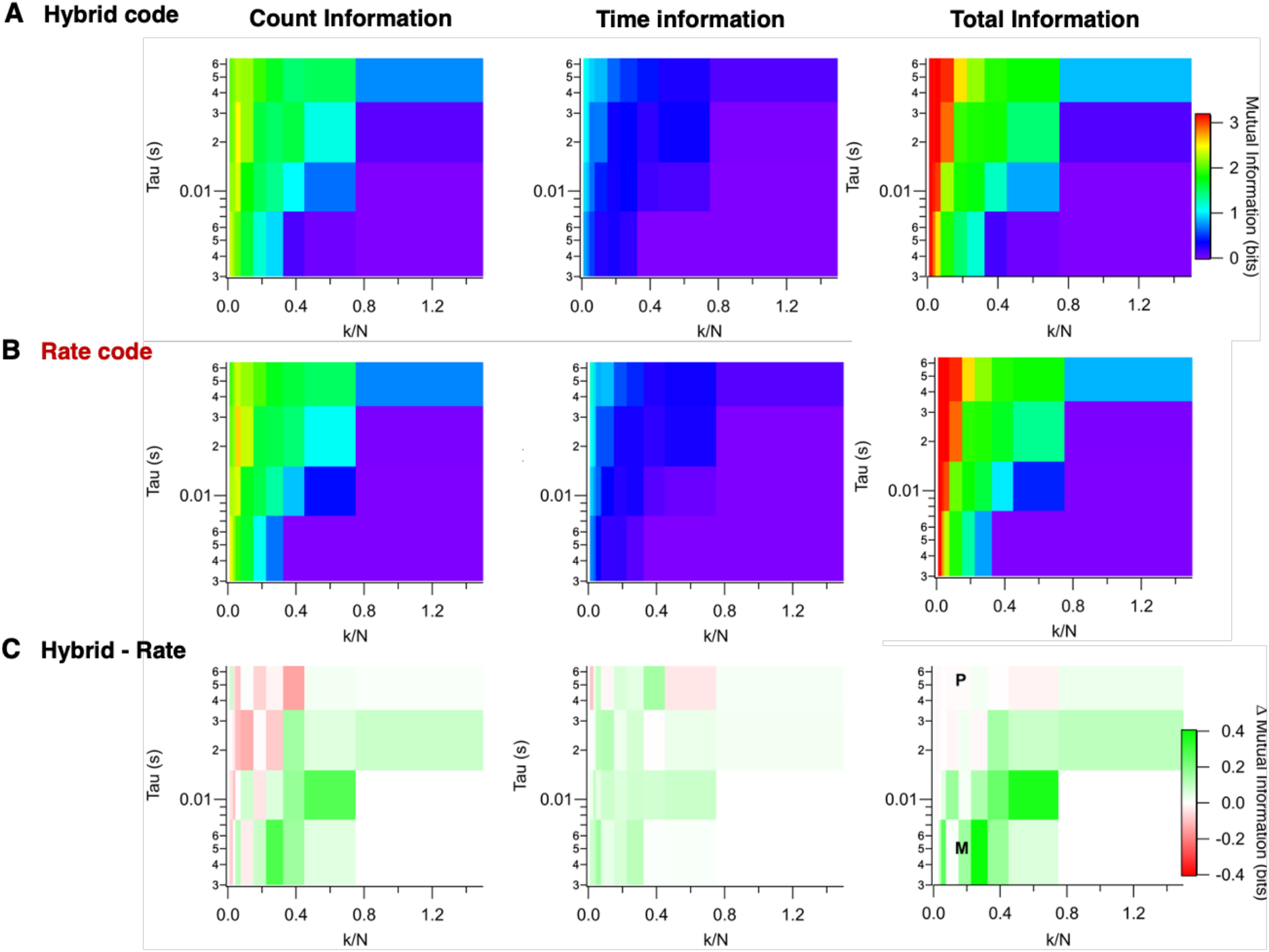
Hybrid coding increased information in the spike output over a range of physiologically-relevant conditions. **A**. Left: heat map showing the information transmitted in the spike count generated by hybrid synaptic inputs for cells of varying k/N and τ_l_. Mutual Information was averaged over values of k and N shown in Table 1 and was measured over a time-window of 200 ms. These values can therefore be multiplied by 5 to obtain information rates as bits s^-1^. Middle: Mutual Information in spike timing. Right: the total Mutual Information representing the sum of count and timing information. The colour scale bar applies to all three maps. Maximum possible information was 3.17 bits. More information was contained in the spike count compared to spike times. **B**. As A, but for rate-coded synaptic inputs operating at the same average release rates. **C**. The difference between A and B (hybrid - rate). The colour scale (right) shows combinations of parameters in which hybrid inputs generated more spikes in green. Zero difference is white and combinations of parameters in which rate-coded inputs generated more spikes are in red. Note that there are some situations in which count information is favoured by rate-coded inputs while time information is favoured by hybrid inputs (and vice versa) but total information is favoured by hybrid inputs over a wider range of conditions (right). The parameters for the specific model cells featured in Fig. 3 are indicated as M (midget) and P (peripheral) in the heat plot for the difference in total information.

The rate code carried marginally more of the total information only in larger cells in which greater numbers of less reliable synapses converge. For the large neuron highlighted in Fig. 3, for instance, we calculated mutual information differences (hybrid – rate) of −0.12 bits for spike count and 0.11 bits for spike time, yielding a total information difference of −0.01 bits. The largest difference in information rate in favour of the rate code was only 0.24 bits s^-1^, occurring when only 10 vesicles were required to generate a spike, with 50 inputs, and a time constant of 50 ms.

Together, the simulations in Figs. 2-6 indicate that synapses which generate MVR can often trigger more spikes per released vesicle compared to a classical Poisson synapse in which all vesicles are released independently. In cells with time-constants less than 15 ms, the tendency for MVR to enhance the transmission of information becomes greater as k/N falls below ∼0.8. These conditions are physiologically relevant. In the peripheral retina, for instance, alpha ganglion cells have membrane time-constants less than 5 ms and ten or fewer quanta released from thousands of synapses are sufficient to generate a spike (Freed and Sterling, 1988; O’Brien et al., 2002). More morphologically compact neurons, such as midget ganglion cells in the fovea of primates, only receive inputs from synapses from a single midget bipolar cell and are likely to have longer time-constants (Sinha et al., 2017).

### MVR increases the efficiency of information transmission across a range of conditions

Neurons can transmit more information by spiking at higher rates but this comes at a relatively large energetic cost (Laughlin, 2001). The release of vesicles and generation of spikes is the major consumer of energy in the brain with one estimate being of the order of ∼24,000 ATP molecules consumed per bit of information transmitted (Attwell and Laughlin, 2001; Harris et al., 2012). The efficiency with which vesicles drive the spike code is therefore likely to have been an evolutionary constraint on the vesicle code itself (Sterling and Laughlin, 2015).

To investigate the potential effects of MVR on the efficiency of information transmission we normalized the measurements of total information summarized in Fig. 6A and B as bits per vesicle and bits per spike, as shown in Fig. 7A and B. Considering the amount of information per vesicle, the relative advantages of hybrid and rate codes can be seen to follow a clear pattern: the more reliably inputs responded (i.e the higher k/N) the broader the range of cell time-constants over which hybrid-coded inputs improved the efficiency with which vesicles transferred information through spikes in the target neuron (Fig. 7A right). The rate-code was only slightly advantageous in neurons acting as the best temporal integrators receiving the least reliably responding inputs. The highest vesicle efficiency using either synaptic code was ∼0.044 bits per vesicle.

**Figure 7.**
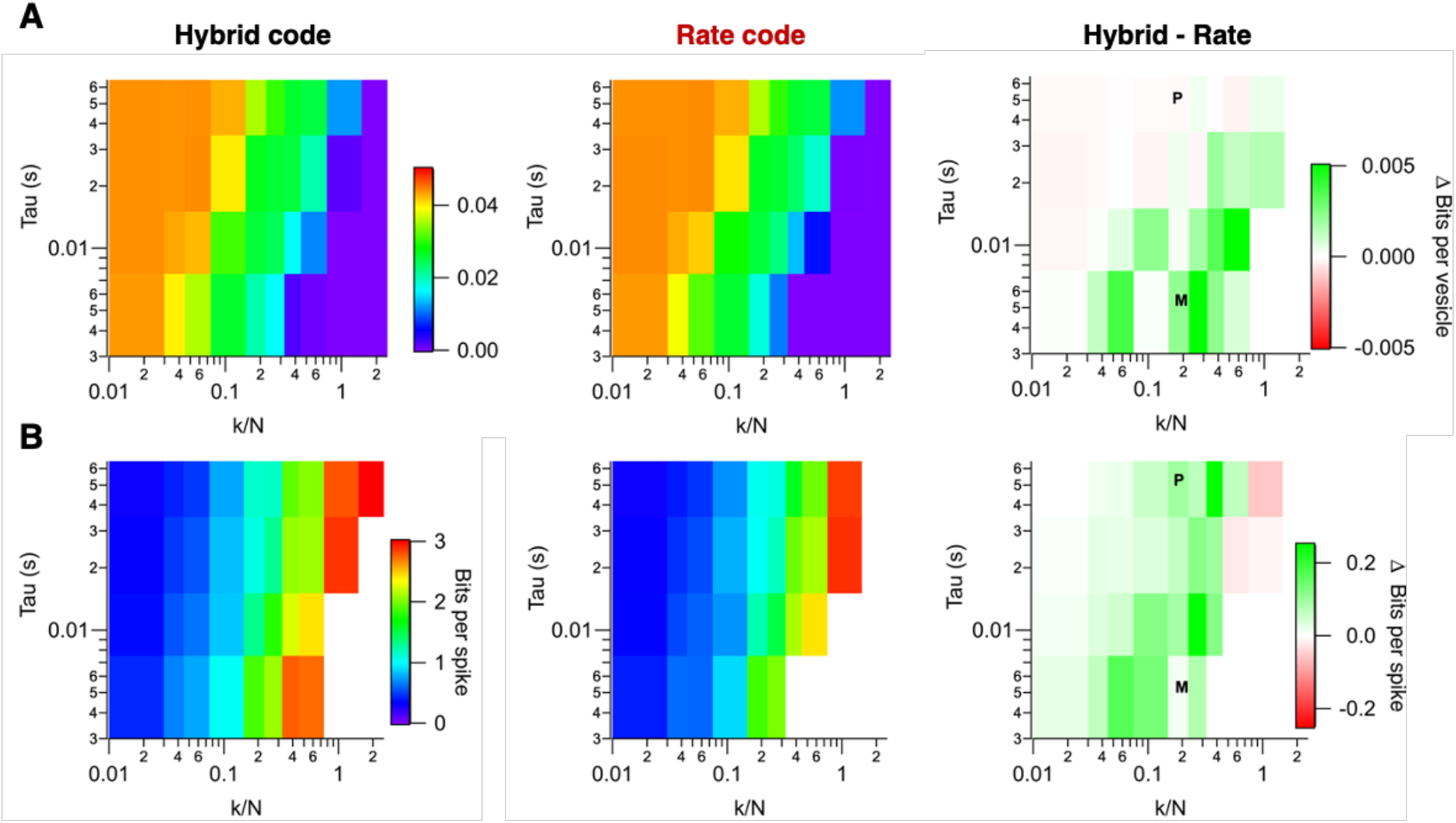
Hybrid coding increased the efficiency of information transmission over a range of conditions. **A**. Left: heat map showing the information transmitted per vesicle generated by hybrid synaptic inputs for cells of varying k/N and τ_l_. Middle: the same for rate-coded synaptic inputs operating at the same average release rates. Right: The difference in the efficiency of information transmission considered as bits per vesicle (hybrid - rate). The colour scale (right) shows combinations of parameters in which hybrid inputs generated more spikes in green. The parameters for the specific model cells featured in Fig. 3 are indicated as M (midget) and P (peripheral) in the heat plot. **B**. As A, but considering efficiency as bits per spike. Note that the hybrid vesicle code increases the efficiency of information transmission across a range of conditions. The pure rate vesicle code was only advantageous in LIF neurons with the longest time-constants driven by the most reliably responding synaptic inputs.

A similar pattern was observed when we quantified the amount of information per spike: the more reliably inputs responded the broader the range of cell time-constants over which hybrid-coded inputs increased the amount of information per spike (Fig. 7B right). The highest spike efficiency was ∼3 bits per spike. A comparison can be made with the information transmitted by spikes in RGCs responding to white-noise stimuli, where the most sluggish transmit ∼3.5 bits/spike, while those that fire most briskly encode ∼2 bits/spike (Koch et al., 2006). The results in Fig. 7 encapsulate what we believe is the most fundamental insight of this study – the potential of MVR to increase the efficiency with which vesicles are used to transmit information.

### The first synapse in hearing

MVR is also a prominent property of the ribbon synapse of auditory hair cells, where the fusion of multiple vesicles can be synchronized on timescales as short as 100 μs (Fuchs and Glowatzki, 2015; Glowatzki and Fuchs, 2002). This presents a special case because there is no convergence onto the target, each Type I afferent fiber receiving sensing glutamate released from just one ribbon synapse. Experiments indicate that spikes in the afferent fiber can be triggered by just a few vesicles and in some cases perhaps as a few as one (Grabner and Moser, 2018; Rutherford et al., 2012). We therefore considered two broad situations: i) k = 1 where *any* vesicular event, regardless of quantal content, is capable of generating a spike, and ii) k > 1, where some (but not all) multivesicular events are capable of generating a spike from rest.

The transmission of information was strongly dependent on both the number of vesicles required to generate a spike, k, and the membrane time constant but again the nature of this dependence differed for synapses employing a pure rate code compared to those also employing MVR (Fig. 8A). Rate coding produced the maximum mutual information in the simulations when k = 1, where the spike sequence in the afferent reproduced the vesicle sequence leaving the synapse (Fig. 7A, red). In this situation a multivesicular event is no more effective than a univesicular event so the hybrid code produces fewer spikes per vesicle and reduces the efficiency of information transmission (Fig. 8A, black).

**Figure 8.**
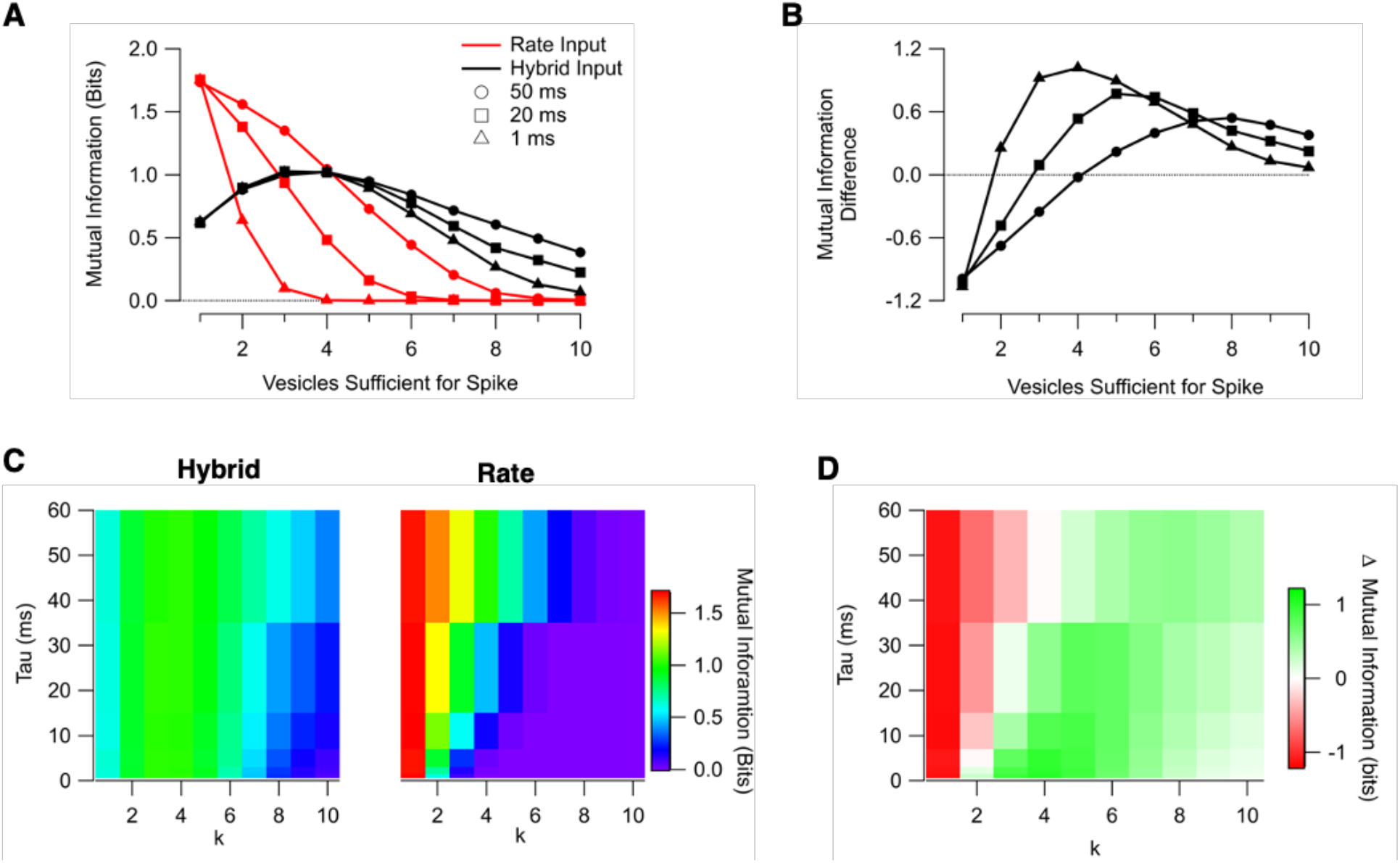
Spike Count Information for Amplitude (black) and Rate (red) inputs as a function of τ_l_ and k where N = 1. **A**. Mutual information for the rate input (dash) and amplitude input (solid) as a function of the vesicles sufficient to generate a spike. Values shown for τ_l_ of 50, 20, 10, and 1 ms. **B**. Mutual information difference (hybrid – rate) for the data shown in A. **C**. Mutual information (as **A**) in matrix form. **D**. Mutual information difference in matrix form.

As the gain of synaptic transmission decreased and more vesicles were required to generate a spike (k > 1), the picture began to change. Increasing k reduced the spike count and effectively abolished spiking at k > 3 when tau = 1 ms (Fig. 8A and B). Essentially, this is because the very short time-constant of the target neuron (mimicking an afferent fiber) did not allow for effective summation of vesicles. Hybrid coding, however, expanded the range of responses by increasing the number of spikes a stimulus could generate, maximizing at a value when most events at higher contrast individually generated a spike (approximately k = 4 for this range of conditions, as seen in Fig. 8B and D). This switch in the relative efficiency of the two synaptic coding strategies starkly illustrates a key functional advantage of the hybrid code: presynaptic integration of vesicles by MVR compensates for situations where the integration time of the post-synaptic neuron is short and limits summation post-synaptically.

The heatmap in Fig. 8D demonstrates that the increase in the efficiency of information transmission caused by MVR was most dramatic in a target neuron with time-constants less than about 10-15 ms in which a spike could be triggered by the arrival of 3-5 vesicles. For 5 < k < 10, MVR still utilized vesicles more efficiently than a pure rate-code (Fig. 8D), despite the fact that not all MVR events were capable of generating a spike even at the highest contrasts. MVR in rat hair cells can generate events equivalent to up to ∼22 vesicles (Glowatzki and Fuchs, 2002) suggesting that the advantages of a hybrid code might be exerted over a broad range of k in the auditory system.

## Discussion

The motivation behind this study was to understand how the unusual statistics of vesicle release from ribbon synapses affected the conversion of analogue sensory signals into spikes. We compared two different regimes of synaptic coding known to operate in different parts of the brain: independent release generating a pure rate code and a hybrid code in which information is contained in both the rate and amplitude of synaptic events. The results indicate that MVR can increase the efficiency with vesicles are used to transmit sensory information over a range of conditions mimicking transmission of visual signals to retinal ganglion cells of different sizes (Figs. 4, 6-7). In the special case of auditory hair cells driving afferent fibers through a single synapse, MVR increased the efficiency of information transfer whenever spike generation required depolarization greater than that caused by a single vesicle (Fig. 8).

Two aspects of the process driving spikes interacted to determine the relative efficiency of a pure rate-code compared to a hybrid code: the time-constant of the target neuron and the reliability of the synaptic inputs. MVR tended to be advantageous when i) the neurons time-constant was shorter, causing less post-synaptic summation of synaptic potentials at physiological rates of vesicle release, and ii) individual synaptic inputs were more reliable so that a lower degree of convergence was required to depolarize the target to threshold for a given stimulus strength.

### What do ribbon synapses employ MVR to transmit sensory signals?

Neurons represent sensory stimuli, at least in part, using a rate-code, where information is contained in the number of spikes over a given time-window (Rieke and Warland, 1999). But it has long been recognized that a potential drawback of a pure rate-code is the time-scale on which it can transmit information: the more variable a neural response to a stimulus the longer the observation time required to detect it to a given degree of certainty (Gerstner et al., 1997; Stein et al., 2005). At the ribbon synapse of bipolar cells, for instance, the maximum sensitivity to changes in contrast occurs around an average release rate R of 20 vesicles s^-1^ and a 10% increase in contrast increases release by a factor of a = 1.8 (James et al., 2019). If release events are Poisson distributed, signal detection theory can be used to quantify the time-window over which the response would need to be observed to differentiate it from ongoing release: a SNR of 2 would require counting an extra 14 vesicles over an observation time (Δt) of ∼0.9 s (equations 7-9 in Methods). But vision operates much more rapidly. Our visual system can process a scene in less than 150 ms (Thorpe et al., 1996) and neurons in the cortex of monkeys can respond to images changing at intervals as short as 14 ms (Keysers et al., 2001).

The convergence of bipolar cell synapses onto RGCs is of course one features of the retinal circuit that will shorten the time-scale of information transmission using a rate-code. N equivalent Poisson inputs allows a given SNR to be achieved with a time-window that varies as 1/√N (Faisal et al., 2008; Gerstner et al., 1997; Rusakov et al., 2020). Continuing with the example above and considering a time-window of 100 ms, the output of 50 synapses would provide a SNR of 2 in a time-window of 100 ms, but this would require the release of N*(1-a)*R*Δt = 80 extra vesicles. This is a very significant cost given that synaptic transmission is a major source of energy consumption in the brain (Attwell and Laughlin, 2001).

The potential problems with a pure rate code highlighted above are closely linked to the idea that the firing rate of a neuron or release rate of a synapse encodes the sensory variable continuously. In fact, if a varying stimulus is applied, most RGCs fire in bursts from rest, driven by fusillades of vesicles released from bipolar cells (Berry et al., 1997b; Freed, 2005; Keat et al., 2001). The information that RGCs convey is therefore represented not just by the spike count but also by the precise timing of spikes (Gutig et al., 2013; Meister et al., 1995; Victor, 2000) and it has been shown that such a temporal coding strategy can significantly improve the speed of information transmission (Butts et al., 2007; Gollisch and Meister, 2008). Crucially, our measurements of glutamate release from individual ribbon synapses of bipolar cells demonstrate that these also signal visual events in bursts, including MVR events of varying amplitude (Fig. 1A and C).

A consideration of different synaptic coding strategies should therefore also consider the representation of temporally discrete events rather than the continuous encoding of a stimulus variable such as contrast. It has been suggested that the onset of a burst in an RGC provides information about when a visual event occurs, while the number of spikes within the burst represent properties such as the magnitude of the event (Keat et al., 2001). The temporal jitter in the first spike within a burst can be as low as a few milliseconds (Berry et al., 1997a; Freed, 2005) and it is notable that larger MVR events from bipolar cells achieve a similar temporal precision (Fig. 1E and F). This comparison highlights one of the potential advantages of representing a stimulus using synaptic symbols of different amplitude: an unexpected symbol immediately imparts new information. In support of this idea, it has been shown that larger MVR events that are more likely to trigger a spike are rarer and carry more information (James et al., 2019).

### Encoding auditory events

The spike code carrying auditory information in cochlear afferents is determined only by excitatory inputs from hair cells and is therefore likely to reflect the temporal precision of ribbon synapses more directly than the spike code leaving the retina. The synaptic output from hair cells has been measured *in vitro* with electrophysiological precision where it is found that larger MVR events are more tightly phase-locked to a sinusoidal voltage-clamp stimulus (Li et al., 2014). Notably, MVR events in hair cells can be as large as ∼20 vesicle equivalents when triggered using electrophysiology (Glowatzki and Fuchs, 2002) but it is unclear if these can also be triggered by sound. Here we used the statistics of MVR measured in bipolar cells *in vivo*, where the largest events evoked by the highest contrasts are composed of 9-11 vesicles (James et al., 2019). A better understanding of how the ribbon synapse of hair cells drive the spike code in auditory fibers will require experimental measurements of the statistics of vesicle release in response to sounds of different intensities and frequencies in hair cells with different temporal tuning.

### Future directions

Our analysis of mutual information was more concerned with the “what” of the stimulus (contrast), and spike count carried more of the total information than spike time for either of the synaptic coding strategies (Fig. 7). This feature of the results might, however, represent limitations of the LIF model in which we only considered excitation. The timing of spikes in RGCs is also dependent on inhibitory inputs, especially feedforward inhibition that increases the temporal precision of spikes and, crucially, also causes spikes to be fired in bursts (Johnston and Lagnado, 2015; Murphy-Baum and Taylor, 2018). Network models incorporating a wider range of biophysical parameters will help us understand how excitation and inhibition interact to determine the temporal precision of the spike code leaving the retina.

To put the statistics of vesicle release into a functional context we also need an understanding of the spike code that it drives. The most decisive next step would be to obtain direct experimental measurements of how effectively coordinated MVR events of different size trigger spikes. The relatively simple organization of the first stage in hearing, where spikes in afferents are driven by a single ribbon synapse, provides a beautiful context in which to ask this question once the experimental tools are established. The situation in the retina will be more complex because RGCs come in a wide variety of morphological and functional types with different time-constants and number and strength of synaptic inputs. In the peripheral retina, for instance, alpha ganglion cells have membrane time-constants less than 5 ms and on average ten quanta are released from thousands of synapses for each spike generated (Freed, 2005). More morphologically compact neurons, such as midget ganglion cells in the fovea of primates, only receive inputs from a single midget bipolar cell and are likely to have much longer time-constants (Sinha et al., 2017). This variety of different functional contexts could also be explored with biophysical models that are more realistic than the LIF neuron that we have begun with here, potentially taking into account non-linearities such as saturation of post-synaptic glutamate receptors and active dendritic conductances.

## Acknowledgements

We thank the Wellcome Trust for funding (221936/Z/20/Z).

## References

Abbott, L.F. (1999). Lapicque’s introduction of the integrate-and-fire model neuron (1907). Brain Res Bull 50, 303–304.

Attwell, D., and Laughlin, S.B. (2001). An energy budget for signaling in the grey matter of the brain. Journal of Cerebral Blood Flow & Metabolism 21, 1133–1145.

Auger, C., Kondo, S., and Marty, A. (1998). Multivesicular release at single functional synaptic sites in cerebellar stellate and basket cells. J Neurosci 18, 4532–4547.

Baden, T., Esposti, F., Nikolaev, A., and Lagnado, L. (2011). Spikes in retinal bipolar cells phase-lock to visual stimuli with millisecond precision. Curr Biol 21, 1859–1869.

Berry, M.J., Warland, D.K., and Meister, M. (1997a). The structure and precision of retinal spike trains. Proceedings of the National Academy of Sciences 94, 5411–5416.

Berry, M.J., Warland, D.K., and Meister, M. (1997b). The structure and precision of retinal spike trains. Proc Natl Acad Sci U S A 94, 5411–5416.

Borst, A., and Theunissen, F.E. (1999). Information theory and neural coding. Nat Neurosci 2, 947–957.

Buracas, G.T., Zador, A.M., DeWeese, M.R., and Albright, T.D. (1998). Efficient discrimination of temporal patterns by motion-sensitive neurons in primate visual cortex. Neuron 20, 959–969.

Burkitt, A.N. (2006a). A review of the integrate-and-fire neuron model: I. Homogeneous synaptic input. Biol Cybern 95, 1–19.

Burkitt, A.N. (2006b). A review of the integrate-and-fire neuron model: II. Inhomogeneous synaptic input and network properties. Biol Cybern 95, 97–112.

Butts, D.A., Weng, C., Jin, J., Alonso, J.M., and Paninski, L. (2011). Temporal precision in the visual pathway through the interplay of excitation and stimulus-driven suppression. J Neurosci 31, 11313–11327.

Butts, D.A., Weng, C., Jin, J., Yeh, C.-I., Lesica, N.A., Alonso, J.-M., and Stanley, G.B. (2007). Temporal precision in the neural code and the timescales of natural vision. Nature 449, 92–95.

Calkins, D.J., Schein, S.J., Tsukamoto, Y., and Sterling, P. (1994). M and L cones in macaque fovea connect to midget ganglion cells by different numbers of excitatory synapses. Nature 371, 70–72.

Choi, S.Y., Borghuis, B.G., Rea, R., Levitan, E.S., Sterling, P., and Kramer, R.H. (2005). Encoding light intensity by the cone photoreceptor synapse. Neuron 48, 555–562.

Christie, J.M., and Jahr, C.E. (2006). Multivesicular release at Schaffer collateral-CA1 hippocampal synapses. J Neurosci 26, 210–216.

Destexhe, A., and Rudolph, M. (2004). Extracting Information from the Power Spectrum of Synaptic Noise. Journal of Computational Neuroscience 17, 327–345.

Faisal, A.A., Selen, L.P., and Wolpert, D.M. (2008). Noise in the nervous system. Nat Rev Neurosci 9, 292–303.

Freed, M.A. (2005). Quantal encoding of information in a retinal ganglion cell. J Neurophysiol 94, 1048–1056.

Freed, M.A., and Sterling, P. (1988). The ON-alpha ganglion cell of the cat retina and its presynaptic cell types. J Neurosci 8, 2303–2320.

Fuchs, P.A., and Glowatzki, E. (2015). Synaptic studies inform the functional diversity of cochlear afferents. Hear Res 330, 18–25.

Gerstner, W., Kreiter, A.K., Markram, H., and Herz, A.V. (1997). Neural codes: firing rates and beyond. Proc Natl Acad Sci U S A 94, 12740–12741.

Glowatzki, E., and Fuchs, P.A. (2002). Transmitter release at the hair cell ribbon synapse. Nat Neurosci 5, 147–154.

Goldberg, J.M., and Brown, P.B. (1969). Response of binaural neurons of dog superior olivary complex to dichotic tonal stimuli: some physiological mechanisms of sound localization. J Neurophysiol 32, 613–636.

Gollisch, T., and Meister, M. (2008). Rapid neural coding in the retina with relative spike latencies. Science 319, 1108–1111.

Gollisch, T., and Meister, M. (2010). Eye smarter than scientists believed: neural computations in circuits of the retina. Neuron 65, 150–164.

Grabner, C.P., and Moser, T. (2018). Individual synaptic vesicles mediate stimulated exocytosis from cochlear inner hair cells. Proc Natl Acad Sci U S A 115, 12811–12816.

Harris, Julia J., Jolivet, R., and Attwell, D. (2012). Synaptic Energy Use and Supply. Neuron 75, 762–777.

Hays, C., Sladek, A., Field, G., and Thoreson, W. (2021). Properties of multi-vesicular release from mouse rod photoreceptors support transmission of single photon responses. eLife 10.

Holler, S., Kostinger, G., Martin, K.A.C., Schuhknecht, G.F.P., and Stratford, K.J. (2021). Structure and function of a neocortical synapse. Nature 591, 111–116.

Huang, C.H., Bao, J., and Sakaba, T. (2010). Multivesicular release differentiates the reliability of synaptic transmission between the visual cortex and the somatosensory cortex. J Neurosci 30, 11994–12004.

Ishii, T., and Hosoya, T. (2020). Interspike intervals within retinal spike bursts combinatorially encode multiple stimulus features. PLoS Comput Biol 16, e1007726.

Jacoby, R.A., Wiechmann, A.F., Amara, S.G., Leighton, B.H., and Marshak, D.W. (2000). Diffuse bipolar cells provide input to OFF parasol ganglion cells in the macaque retina. J Comp Neurol 416, 6–18.

James, B., Darnet, L., Moya-Diaz, J., Seibel, S.H., and Lagnado, L. (2019). An amplitude code transmits information at a visual synapse. Nat Neurosci 22, 1140–1147.

Johnston, J., and Lagnado, L. (2015). General features of the retinal connectome determine the computation of motion anticipation. eLife 4, e06250.

Kaplan, E., Purpura, K., and Shapley, R.M. (1987). Contrast affects the transmission of visual information through the mammalian lateral geniculate nucleus. The Journal of physiology 391, 267–288.

Kayser, C., Montemurro, M.A., Logothetis, N.K., and Panzeri, S. (2009). Spike-Phase Coding Boosts and Stabilizes Information Carried by Spatial and Temporal Spike Patterns. Neuron 61, 597–608.

Keat, J., Reinagel, P., Reid, R.C., and Meister, M. (2001). Predicting Every Spike: A Model for the Responses of Visual Neurons. Neuron 30, 803–817.

Keysers, C., Xiao, D.-K., Földiák, P., and Perrett, D.I. (2001). The speed of sight. Journal of cognitive neuroscience 13, 90–101.

Koch, K., McLean, J., Segev, R., Freed, M.A., Berry, M.J., 2nd, Balasubramanian, V., and Sterling, P. (2006). How much the eye tells the brain. Curr Biol 16, 1428–1434.

Kolb, H., and Marshak, D. (2003). The midget pathways of the primate retina. Doc Ophthalmol 106, 67–81.

Lagnado, L., and Schmitz, F. (2015). Ribbon Synapses and Visual Processing in the Retina. Annu Rev Vis Sci 1, 235–262.

Laughlin, S.B. (2001). Energy as a constraint on the coding and processing of sensory information. Curr Opin Neurobiol 11, 475–480.

Levine, M.W. (1991). The distribution of the intervals between neural impulses in the maintained discharges of retinal ganglion cells. Biological Cybernetics 65, 459–467.

Li, G.L., Cho, S., and von Gersdorff, H. (2014). Phase-locking precision is enhanced by multiquantal release at an auditory hair cell ribbon synapse. Neuron 83, 1404–1417.

Li, X., and Ascoli, G.A. (2008). Effects of synaptic synchrony on the neuronal input-output relationship. Neural Comput 20, 1717–1731.

Lisman, J.E., Raghavachari, S., and Tsien, R.W. (2007). The sequence of events that underlie quantal transmission at central glutamatergic synapses. Nat Rev Neurosci 8, 597–609.

Meister, M., and Berry, M.J., 2nd (1999). The neural code of the retina. Neuron 22, 435–450.

Moya-Díaz, J., James, B., Esposti, F., Johnston, J., and Lagnado, L. (2022). Diurnal changes in the efficiency of information transmission at a sensory synapse. bioRxiv, 2021.2009.2012.459944.

Murphy-Baum, B.L., and Taylor, W.R. (2018). Diverse inhibitory and excitatory mechanisms shape temporal tuning in transient OFF α ganglion cells in the rabbit retina. The Journal of physiology 596, 477–495.

Neef, A., Khimich, D., Pirih, P., Riedel, D., Wolf, F., and Moser, T. (2007). Probing the mechanism of exocytosis at the hair cell ribbon synapse. J Neurosci 27, 12933–12944.

O’Brien, B.J., Isayama, T., Richardson, R., and Berson, D.M. (2002). Intrinsic physiological properties of cat retinal ganglion cells. J Physiol 538, 787–802.

Odermatt, B., Nikolaev, A., and Lagnado, L. (2012). Encoding of luminance and contrast by linear and nonlinear synapses in the retina. Neuron 73, 758–773.

Panzeri, S., Senatore, R., Montemurro, M.A., and Petersen, R.S. (2007). Correcting for the sampling bias problem in spike train information measures. J Neurophysiol 98, 1064–1072.

Pillow, J.W., Paninski, L., Uzzell, V.J., Simoncelli, E.P., and Chichilnisky, E.J. (2005). Prediction and decoding of retinal ganglion cell responses with a probabilistic spiking model. J Neurosci 25, 11003–11013.

Rathbun, D.L., Warland, D.K., and Usrey, W.M. (2010). Spike timing and information transmission at retinogeniculate synapses. J Neurosci 30, 13558–13566.

Reinagel, P., and Reid, R.C. (2002). Precise firing events are conserved across neurons. J Neurosci 22, 6837–6841.

Rieke, F., and Warland, D. (1999). Spikes: exploring the neural code (MIT press).

Rusakov, D.A., Savtchenko, L.P., and Latham, P.E. (2020). Noisy Synaptic Conductance: Bug or a Feature? Trends Neurosci 43, 363–372.

Rutherford, M.A., Chapochnikov, N.M., and Moser, T. (2012). Spike encoding of neurotransmitter release timing by spiral ganglion neurons of the cochlea. J Neurosci 32, 4773–4789.

Shannon, C. (1948). A Mathematical Theory of Communication. Bell System Technical Journal 27, 379–423.

Singer, J.H., Lassova, L., Vardi, N., and Diamond, J.S. (2004). Coordinated multivesicular release at a mammalian ribbon synapse. Nat Neurosci 7, 826–833.

Sinha, R., Hoon, M., Baudin, J., Okawa, H., Wong, R.O.L., and Rieke, F. (2017). Cellular and Circuit Mechanisms Shaping the Perceptual Properties of the Primate Fovea. Cell 168, 413-426.e412.

Stein, R.B., Gossen, E.R., and Jones, K.E. (2005). Neuronal variability: noise or part of the signal? Nature Reviews Neuroscience 6, 389–397.

Sterling, P., and Laughlin, S. (2015). Principles of neural design (MIT press).

Thorpe, S., Fize, D., and Marlot, C. (1996). Speed of processing in the human visual system. Nature 381, 520–522.

Usrey, W.M., and Reid, R.C. (1999). Synchronous activity in the visual system. Annual review of physiology 61, 435–456.

Usrey, W.M., Reppas, J.B., and Reid, R.C. (1998). Paired-spike interactions and synaptic efficacy of retinal inputs to the thalamus. Nature 395, 384–387.

Uzzell, V.J., and Chichilnisky, E.J. (2004). Precision of spike trains in primate retinal ganglion cells. J Neurophysiol 92, 780–789.

Victor, J.D. (2006). Approaches to Information-Theoretic Analysis of Neural Activity. Biol Theory 1, 302–316.

